# Persistent parental RNAi in the beetle *Tribolium castaneum* involves maternal transmission of long double-stranded RNA

**DOI:** 10.1101/2021.11.12.468425

**Authors:** Thorsten Horn, Kalin D. Narov, Kristen A. Panfilio

## Abstract

Parental RNA interference (pRNAi) is a powerful and widely used method for gene-specific knockdown. Yet in insects its efficacy varies between species, and how the systemic RNAi response is transmitted from mother to offspring remains elusive. Using the flour beetle *Tribolium castaneum*, we report an RT-qPCR strategy to unmask the presence of double-stranded RNA (dsRNA) distinct from endogenous mRNA. We find that the injected dsRNA is directly transmitted into the egg and persists throughout embryogenesis. Despite this depletion of dsRNA from the mother, we show that strong pRNAi can persist for months before waning at strain-specific rates. In seeking the receptor proteins for cellular uptake of long dsRNA into the egg, we lastly present a phylogenomics profiling approach to ascertain macroevolutionary distributions of candidate proteins. We demonstrate a visualization strategy based on taxonomically hierarchical assessment of orthology clustering data to rapidly assess gene age and copy number changes, refined by several lines of sequence-based evidence. We use this approach to document repeated losses of SID-1-like channel proteins in the arthropods, including wholesale loss in the Heteroptera (true bugs), which are nonetheless highly sensitive to pRNAi. Overall, we elucidate practical considerations for insect pRNAi against a backdrop of outstanding questions on the molecular mechanism of dsRNA transmission to achieve long-term, systemic knockdown.

## INTRODUCTION

Since the demonstration of systemic RNA interference in insects about twenty years ago [1-3], this technique has become widely used for genetics research and there is growing interest in its application for species- and gene-specific pest management [4-9]. In many species, systemic knockdown is efficient across life history stages, with a particular advantage of parental RNAi. Delivery of dsRNA into the mother, often by a single injection, can achieve knockdown of both maternal and zygotic gene expression in offspring, including at postembryonic stages [10]. This technique can provide highly efficient gene knockdown in hundreds of embryos that are often collected for up to three weeks after injection (*e*.*g*., [1, 11]).

As a well-established model system, the red flour beetle *Tribolium castaneum* has been at the forefront of research on the RNAi mechanism [1, 10, 12] and for diverse genetics studies [13]. It is an effective RNAi screening platform [14-16]. pRNAi in *Tribolium* is regularly used for phenotypic investigation of development and to test genetic interactions singly or globally, such as by RNA-seq after RNAi [17-20]. Empirical work has shown that efficient RNAi is achieved through the introduction of long dsRNA into the organism, which persists longer *in vivo* and has more efficient cellular uptake than short interfering RNA (siRNA) [10, 21]. Supporting this, an early genomic survey of RNAi molecular machinery in *Tribolium* [12] confirmed conservation of many core elements, but also with notable absences or changes in copy number or function of some elements compared to the well understood RNAi system of *C. elegans*. This has generally been borne out by studies in other insect species [4, 5].

However, the mechanism of pRNAi is still poorly understood. Germline tissues and developing eggs have been studied as one of several tissue types that exhibit distinct susceptibilities to systemic knockdown in adult females. On the one hand, germline tissue showed lower levels of systemic effect in a pea aphid study in which this tissue was distal to the site of initial dsRNA delivery [9]. On the other hand, research in *C. elegans* has shown co-localization of dsRNA and yolk in oocytes, suggesting dsRNA transmission via a general mechanism for maternal provisioning of eggs [22].

A key element for elucidating systemic pRNAi is the ability to detect and track the dsRNA. In *C. elegans*, microscopy for visual detection of fluorescently labeled dsRNA showed that 50-bp dsRNA was transmitted to the oocyte [22]. However, this qualitative study did not examine embryos beyond the four-cell stage or test long dsRNA (∼400 bp for efficient knockdown in *Tribolium*, [10, 15]). Visual tracking of fluorescently labeled dsRNA has been attempted in insects, but with limits on transmissibility and detection sensitivity [23, 24]. Recent reviews on insect RNAi have thus explicitly called for the use of quantitative, sensitive detection methods such as RT-qPCR as a complementary approach: both to assay the extent of target gene knockdown after RNAi and for the systematic tracking of dsRNA [6]. RT-qPCR to assay knockdown is regularly used in developmental genetics research [19, 20, 25], as one of several methods alongside global assays such as RNA-seq [17, 20] and spatiotemporally sensitive methods such as *in situ* hybridization, which can also detect inter-embryo variability (*e*.*g*., [25]). To the best of our knowledge, these methods have thus far been used to measure expression levels of endogenous target gene mRNA, but not for dsRNA detection.

Here, we combine experimental results in *Tribolium* with comparative genomics assessments of gene repertoires across species to shed further light on the molecular mechanisms of dsRNA transmission during systemic pRNAi in insects. We present an RT-qPCR strategy whose amplicon design and sensitivity distinguishes dsRNA in offspring after pRNAi for genes with distinct temporal expression profiles, demonstrating its value for tracking throughout embryogenesis. Furthermore, we show that knockdown in progeny persists at high levels for months, despite a finite starting amount of dsRNA, through time-course analyses that evaluate female age, genetic strain, and different target genes. Lastly, we compare hundreds of sequenced animal genomes to reveal limits in the conservation of candidate receptor proteins for dsRNA uptake, emphasizing the specificity of the importer protein SID-1 to nematodes compared to insects or vertebrates. Thus, even as we provide empirical advances for investigation and application of pRNAi, we also flag multiple aspects of dsRNA transport that remain enigmatic.

## RESULTS

### dsRNA is transported into eggs and persists during embryogenesis

The homeodomain transcription factor Tc-Zen1 is a critical regulator in early development, specifying the identity of the extraembryonic serosal tissue that surrounds the embryo and confers mechanical, physiological, and immunological protection [18, 20, 26, 27]. During routine verification of *Tc-zen1* parental RNAi using RT-qPCR (as in [20]), we unexpectedly found that measured expression of *Tc-zen1* mRNA was higher in RNAi samples than in wild type under certain assay conditions, despite strong phenotypic validation of systemic knockdown (see Methods).

We observed this effect when using an RT-qPCR amplicon that was designed to be small and intron-spanning, ensuring efficient and specific amplification [28, 29]. However, due to the small size of the *Tc-zen1* mRNA transcript, this amplicon was also nested within the region used as an established multi-purpose template for dsRNA and *in situ* hybridization (Fig. 1A: Fragment 2, compared to the long dsRNA, [20, 25, 30]). Using this amplicon, at young embryonic stages we observed strong reduction to 25% of wild type levels in the RNAi sample, consistent with our phenotypic validation (Fig. 1B at 8-24 h: mean expression ratios of 1.24 RNAi/ 4.88 wild type for Fragment 2). In contrast, this amplicon produces higher expression estimates in RNAi than in wild type samples at the older stages assayed (Fig. 1B: yellow *vs*. red plot lines, developmental time ≥16-24 h). When the same samples are assayed with an RT-qPCR amplicon that only partially overlaps the dsRNA fragment (Fig. 1A: Fragment 1), we obtain the expected result of strong RNAi knockdown at all stages, including to only 5% of wild type levels at 8-24 h (mean expression ratios of 0.22 RNAi / 4.51 wild type), and no ostensible overexpression at older stages (Fig. 1B: blue plot lines).

**Figure 1.**
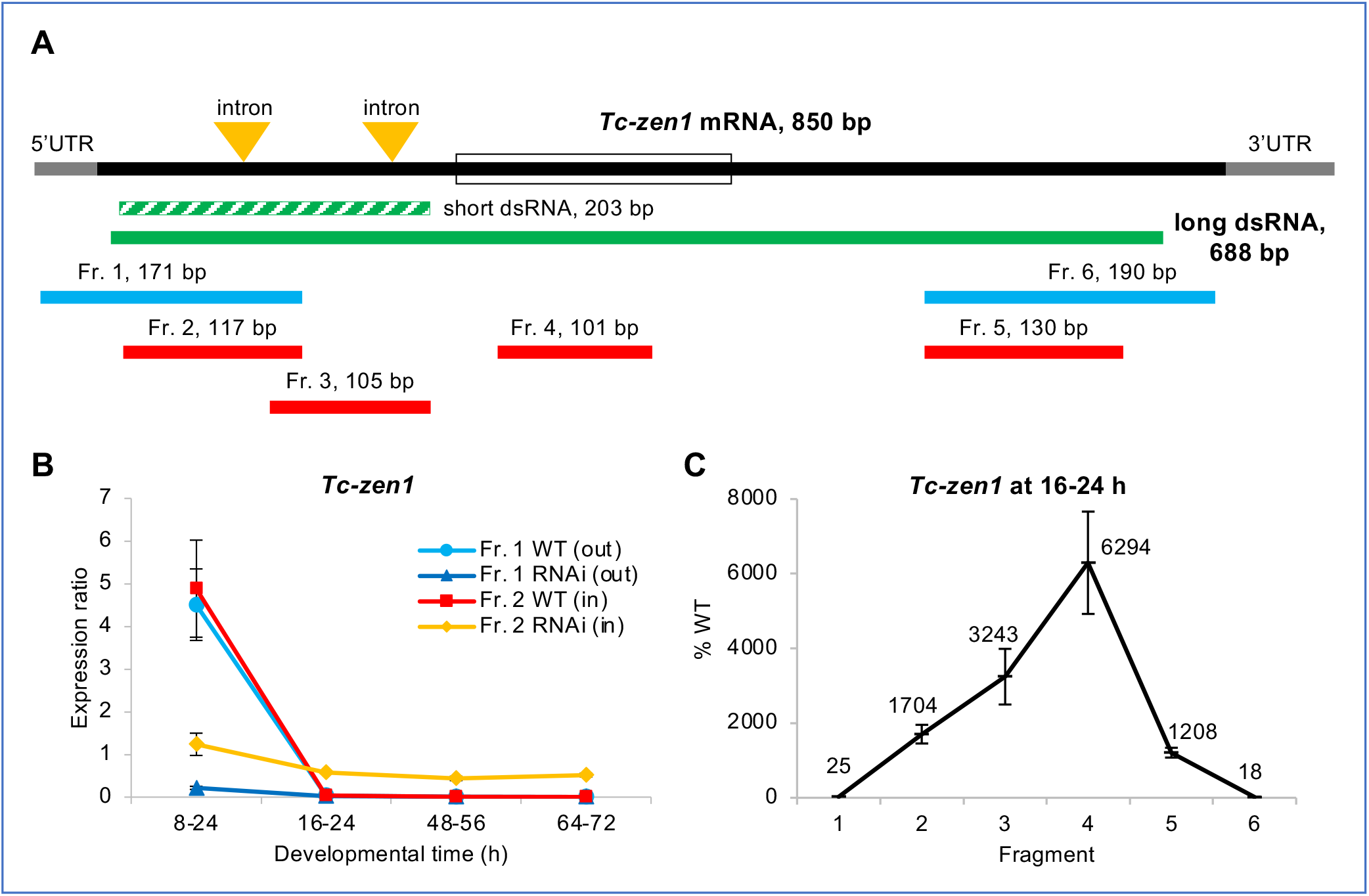
Long dsRNA molecules are transmitted maternally and persist throughout embryogenesis after parental RNAi for *Tc-zen1*. **(A)** Structure of *Tc-zen1* mRNA (CDS: solid black, UTRs: grey, homeobox: open box) and corresponding dsRNA fragments (green) used to silence the gene: the long dsRNA (solid green) was used in this study; the short dsRNA (dashed green) was used previously [20] to specifically avoid the highly conserved homeobox. Beneath, the six fragments (Fr. 1-6) indicate the regions used for RT-qPCR quantification, where the two outermost fragments (blue) lay partially outside of the dsRNA fragment and four fragments (red) lay inside the dsRNA fragment. Fragment lengths are indicated and are shown to scale. **(B)** Expression ratio of *Tc-zen1* in knockdown (RNAi) and wild type (WT) samples at different stages of development, assayed by RT-qPCR with fragments that extend outside (Fr. 1) or are nested within (Fr. 2) the dsRNA fragment, as indicated in the legend. In the three older stages, Fragment 2 in the RNAi samples (yellow) shows consistently higher expression than all other samples, due to its ability to detect the dsRNA in addition to endogenous transcript. Developmental time is specified in hours after egg lay (*i*.*e*., after fertilization). **(C)** *Tc-zen1* expression measured by RT-qPCR in the RNAi samples compared to WT samples for all fragments, at a developmental stage when endogenous mRNA levels are negligible (at 16-24 h). The two outermost fragments (1 and 6) show reduced expression compared to WT, consistent with successful RNAi knockdown, while the inner fragments (2-5) show increased expression after RNAi, with highest overexpression for Fragment 4 (see also Fig. S2). The mean values (%) for each fragment are indicated. Mean expression levels are shown from three biological replicates (see Methods); error bars represent ± one standard deviation.

Notably, the semi-nested amplicon detects the same levels of wild type expression as in our original assay (Fig. 1B: light blue and red plot lines, respectively). This corroborates the accuracy of the original, nested amplicon for quantification of *Tc-zen1* transcript levels. Moreover, these findings with either amplicon are consistent with our previous work that documented a single early pulse of *Tc-zen1* expression that peaks at 6-10 h before rapidly declining to undetectable levels for the rest of embryogenesis [20].

Thus, we infer that after *Tc-zen1* RNAi the nested RT-qPCR amplicon is detecting both residual endogenous transcript as well as dsRNA transmitted from the mother to the egg. This implies that the ostensible overexpression represents the unmasked detection of dsRNA specifically at older developmental stages when wild type expression is low. Under standard culturing conditions, *Tribolium* embryogenesis is about three days, and here we show that the transmitted dsRNA stably persists in the egg throughout this interval (Fig. 1B: yellow plot line, ≥16-24 h). Furthermore, although the nested fragment did capture the reduction in the target gene at a stage of high endogenous expression (8-24 h), the degree of transcript depletion after RNAi is likely underestimated due to the detection of the dsRNA (reduction to 25% with nested Fragment 2 *vs*. to 5% with semi-nested Fragment 1). In summary, there is a certain amount of dsRNA transmitted from the mother to the offspring that is detectable by RT-qPCR, but at levels that may be masked by high endogenous expression.

### The entire long dsRNA molecule is maternally transmitted

The RNAi pathway involves processing of long dsRNA by the RNase III endonuclease Dicer to generate siRNAs of ∼20-23 bp, which is the means of amplifying the RNAi effect to systemic levels [31]. Yet, our nested RT-qPCR amplicon is >100 bp. We thus considered the possibility that the dsRNA is transmitted from the injected mother to the embryo as a largely intact, unprocessed molecule.

Our method to detect transmitted dsRNA relies on measuring different expression levels in the same sample with two different amplicons, one being partially outside of the dsRNA sequence. In theory, this method could also be used to determine the size of the transmitted dsRNA by increasing the length of the amplicons (*e*.*g*., by extending Fragments 1 and 2 in the 3’ direction). Unfortunately, RT-qPCR analysis becomes increasingly unreliable with increasing amplicon size [29], and our results were inconclusive between biological and technical replicates with this strategy.

As an alternative approach, we could robustly measure the relative expression of a series of RT-qPCR amplicons that span the *Tc-zen1* transcript (Fig. 1A: Fragments 1-6). As wild type expression is negligible at 16-24 hours (Fig. 1B), the measured expression at this stage largely represents transmitted dsRNA present in the egg. Validating RNAi efficiency, the two amplicons that lay partially outside the dsRNA region show efficient knockdown of *Tc-zen1* at 16-24 hours (Fig. 1C: Fragments 1 and 6, mean reductions to ≤25% of WT levels). This is consistent with phenotypic validation and RT-qPCR assays of early developmental samples with high wild type expression (Fig. 1B: 8-24 h). In contrast, all amplicons that were fully nested within the dsRNA region show substantially increased expression after RNAi (>1000%; Fig. 1C: Fragments 2-5). Strikingly, there was a five-fold range in expression levels among the nested amplicons, an issue we address in the Discussion in terms of experimental design and gene-specific sequence features. Regardless, these four amplicons are each >100 bp and together span 654 bp. We thus conclude that the entire 688-bp dsRNA molecule injected into the mother is transmitted to the egg.

### Unmasked dsRNA presence at stages of low expression is a general feature

We next expanded our analyses to test whether maternal dsRNA transmission is a general feature of systemic RNAi in *Tribolium*. For this purpose, we chose two additional genes that have distinct, well-characterized expression time courses and molecular functions that differ from *Tc-zen1* and from one another. The first gene, *Tc-chitin synthase 1* (*Tc-chs1*), encodes a large, transmembrane enzyme that extrudes the polysaccharide chitin into developing cuticle of the serosa (early embryogenesis, [27]) and of the larval epidermis (late embryogenesis, [32]). Secondly, in the nuclear GFP (nGFP) line [33], red fluorescence encoded by *DsRed* serves as a transgenic marker under the control of the synthetic Pax6 core promoter-enhancer element 3xP3, which drives late expression in the developing eyes and ventral nerve cord (central nervous system, [34, 35]).

For both genes we detected greater expression in the RNAi samples with the nested amplicon compared to the semi-nested amplicon (Fig. 2A-B: yellow *vs*. dark blue plot lines). Furthermore, the effect was again most pronounced – with ostensible overexpression – at developmental stages when wild type expression is low: early embryogenesis for *DsRed* (4733%) and mid-embryogenesis for *Tc-chs1* (322%). As we had observed this effect in late embryogenesis for *Tc-zen1* (Fig. 1B), these results clarify that it is the level of endogenous expression, and not a specific developmental stage, that determines when dsRNA transmission can be unmasked by our RT-qPCR strategy. This is applicable whether the gene has a single stage of peak expression (*Tc-zen1, DsRed*) or a bimodal temporal expression profile with only a transient period of low expression (*Tc-chs1*). At stages when the target gene is moderately to strongly expressed, for both *Tc-chs1* and 3xP3-driven *DsRed* the nested amplicon underestimates the level of knockdown after RNAi by 5-20%, similar to what we had observed for *Tc-zen1*.

**Figure 2.**
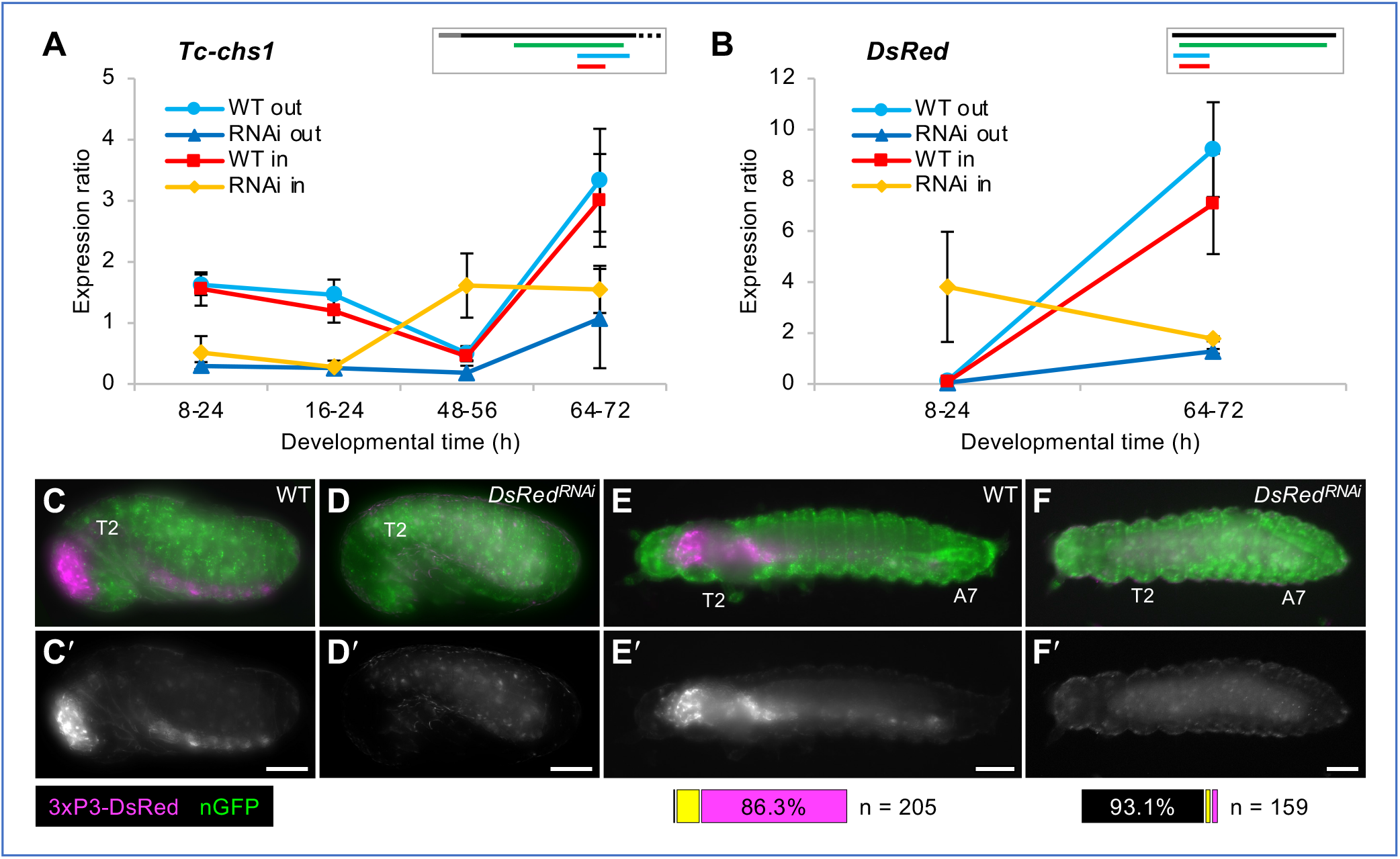
Maternal transmission of dsRNA occurs for diverse genes with distinct expression profiles. **(A-B)** RT-qPCR expression ratio assayed with amplicons that are nested (“in”: red and yellow) or partially outside (“out”: light and dark blue) with respect to the dsRNA fragment, in WT and after RNAi, as indicated in the legends. Mean expression levels are shown from three biological replicates; error bars represent ± one standard deviation. For *Tc-chs1* (A), the nested qPCR amplicon shows higher expression in RNAi samples (yellow) when endogenous *Tc-chs1* expression is low (48-56 h). Similarly, in the nGFP strain expressing transgenic dsRed (B), the *DsRed* nested qPCR amplicon detects a relative overexpression after RNAi at a stage when *DsRed* transgene is not expressed (8-24 h). Inset schematics depict the transcript, dsRNA, and qPCR fragments to scale, using the same color scheme as in Fig. 1; only the first 700 bp of the 5092-bp mRNA is shown for *Tc-chs1*. **(C-F)** Phenotypic confirmation of *DsRed* knockdown through loss of DsRed fluorescence in a transgenic line that ubiquitously expresses nuclear-localized GFP (green). The 3xP3 core promoter drives DsRed signal (magenta) in the brain and ventral nerve cord of untreated control (WT) embryos (C) and larvae (E). After *DsRed* RNAi, 3xP3-driven DsRed signal is absent, with only weak autofluorescence detected in the epidermal cuticle and the yolk (D, F). Views are lateral (C-D) or dorsal (E-F), with anterior left and, as applicable, dorsal up. Landmark thoracic (T) and abdominal (A) segments are numbered. Letter-prime panels show the DsRed channel alone. Scale bars are 100 µm. Horizontal bar charts show the proportions of larvae with no (black), weak (yellow), or strong (magenta) DsRed signal in larvae.

We also verified the knockdown efficiency for *DsRed* in the nGFP line by observing red fluorescence in late embryos and young larvae (Fig. 2C-F). Fluorescent signal was detectable in >99% of untreated (wild type) larvae (n= 205) and absent in 93.1% of RNAi larvae (n= 159), consistent with very high efficiency knockdown.

### pRNAi is highly efficient for months before waning at strain-specific rates

A single injection of the mother provides a finite number of dsRNA molecules, and the knockdown effect of pRNAi wanes over time in insects [1, 3, 36]. Our results suggest that waning may reflect not only endogenous transcript recovery after dsRNA degradation in the mother, but also maternal depletion of dsRNA due to its direct transmission into offspring. To determine how long pRNAi knockdown persists in *Tribolium*, we conducted time course experiments until the knockdown effect had fully waned, testing different genes, genetic backgrounds, and ages of adult female. For this purpose, larval cuticle preparations were used as a robust phenotype assay (see Methods), targeting two genes whose knockdown produces distinctive and easily scorable cuticle phenotypes with high penetrance (Fig. 3A-C): *Tc-tailup* (*Tc-tup*, [15, 37, 38]) and *Tc-germ cell-less* (*Tc-gcl*, [39]).

**Figure 3.**
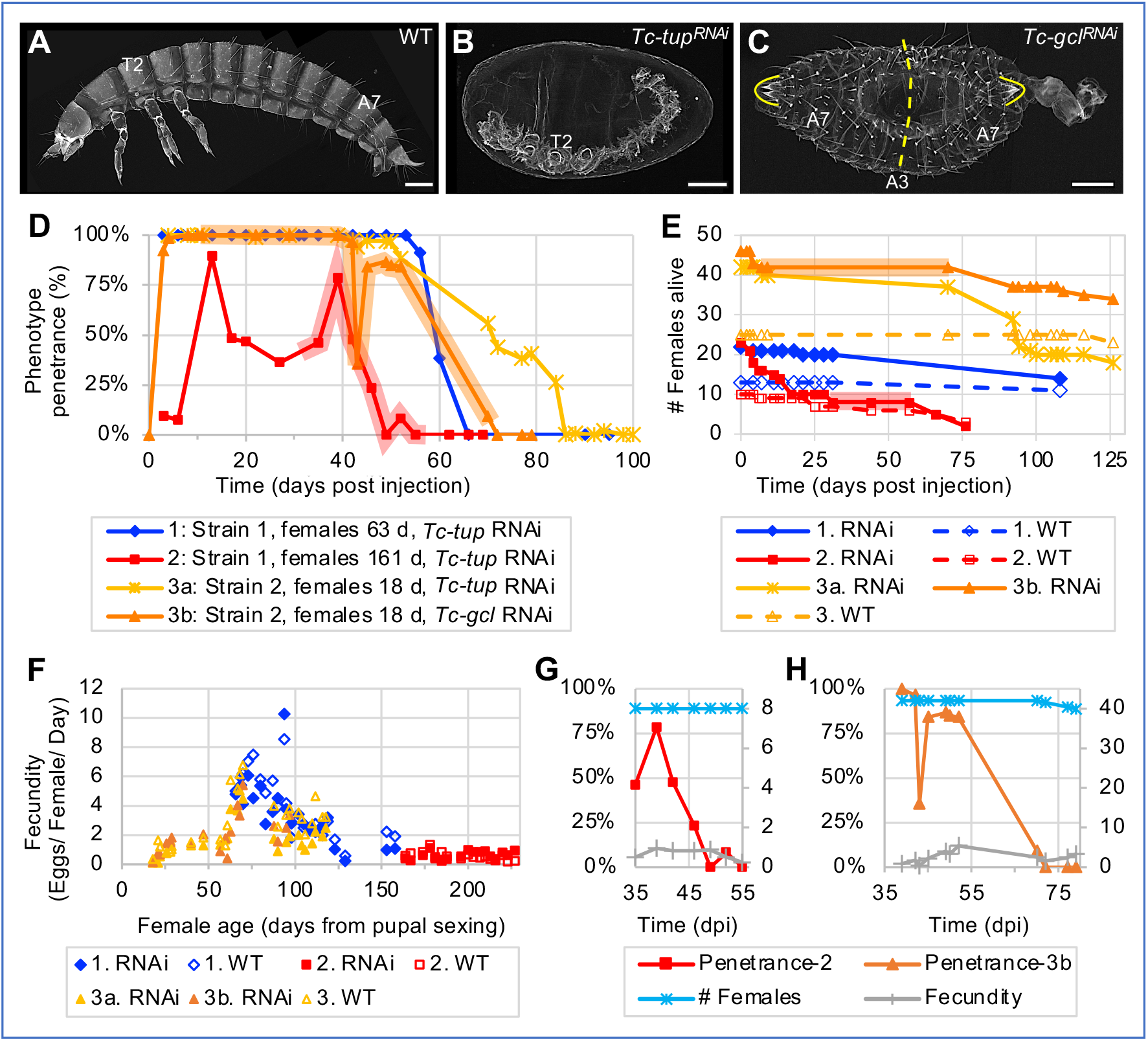
Systemic parental RNAi persists at high levels for months before fully waning. **(A-C)** Representative larval cuticle preparations for wild type (WT), *Tc-tup*^*RNAi*^, and *Tc-gcl*^*RNAi*^ (from Experiment 3, collected 39-52 dpi, assayed ≥6 days after egg lay). Views are lateral (A,B) or dorsal-lateral (C), with anterior left and dorsal up. Landmark thoracic (T) and abdominal (A) segments are numbered. The dashed line indicates the plane of symmetry in the *Tc-gcl*^*RNAi*^ mirror-image double abdomen phenotype; brackets outline the terminal urogomphi. Scale bars are 100 µm. **(D)** Time courses of parental RNAi penetrance from experiments that differ in beetle strain, female age, and target gene for knockdown (see figure legend and Methods). Data points represent minimum age after injection, with n≥10 eggs in each sample (see Methods). Shaded plot segments for Experiments 2 and 3b represent time intervals with dynamic changes in RNAi penetrance that encompass both transient fluctuations (increase or decrease) and the interval of RNAi waning, while female population size was constant (no fatalities). **(E)** Survival curves for females from all treatment conditions from all three experiments. For Experiments 2 and 3b, respectively, the red and orange shading corresponds to the same intervals as in (A). **(F)** Fecundity values (number of eggs per female per day) relative to female age from all treatment conditions in all experiments, assayed at 19-26 time points per treatment. **(G-H)** Juxtaposition of phenotype penetrance (%, left y-axis) with female population size and fecundity values (integer values, right y-axis) for the period of RNAi waning in Experiments 2 and 3b (red and orange shaded intervals, as above): female population size and fecundity remain steady or exhibit only minor fluctuation while RNAi wanes.

Across beetle strains and target genes, >90% penetrance for gene-specific knockdown in embryos is achieved within three days after adult injection and remains persistently high for nearly two months at 30 °C (Fig. 3D: Experiments 1, 3a, and 3b). Only in our aged female experiment did we see a delay in onset of knockdown and lower overall levels of penetrance (generally 50% over a 30-day interval; Fig. 3D: Experiment 2). Nonetheless, across all experiments we still observed 50% phenotype penetrance at 42-71 days after injection. A minor resurgence (<10%) after full depletion of the RNAi phenotype occurred briefly towards the end of both Experiments 2 and 3a.

In contrast to the consistent duration of strong knockdown, the rate of waning may be strain-specific, irrespective of female age or target gene. In Strain 1, knockdown fully declined in a 10-day interval (from 91% or 78% to 0% in Experiments 1 and 2, respectively). Waning in Strain 2 was more gradual, spanning the better part of a month (from ∼86% to 0% over 20-34 days in Experiments 3a and 3b).

### pRNAi waning and transient fluctuations are strain- and female-specific

Since our experimental beetle populations were maintained as pooled cohorts, we examined female lethality and fecundity to more precisely document the pRNAi waning effect (Fig. 3E-H).

Regarding survival (Fig. 3E), the dsRNA-injected females exhibited minor fatalities within the first week after injection before the populations stabilized over the next 1-2 months, until death occurred from presumed old age. The exception to this trend was in Experiment 2, where females were already aged for 5.3 months as adults before injection and subsequent mating: these injected females showed steady mortality for the first 2.5 weeks before the population stabilized through the second month of the experiment. Fatalities of the uninjected (wild type) females and males were minimal in all experiments.

We then determined fecundity in terms of egg output per female per day (Fig. 3F). Age is the strongest predictor of female fecundity; neither the background genetic strain nor dsRNA injection had an appreciable effect. Fecundity fluctuates on short time scales (<1 week), but overall we find a marked but inexplicable increase in fecundity at 50-75 days, with ≥6 eggs/female/ day. After, there is a rapid decline to 130 days, and persistent, low-level fecundity through 230 days.

In sum, we find that on multi-month timescales both survival and egg output of RNAi females is comparable to that of the uninjected controls, indicating that long-term activity of RNAi machinery does not generally impair female physiology or fecundity.

Arguably, intermediate RNAi penetrance at the population level could reflect offspring contributions from a mix of females with strong RNAi and resistant females that only lay wild type offspring. Then, waning of RNAi over time might reflect the earlier death of the females that produced affected offspring. However, our data support the waning of RNAi in individual females. Firstly, for months we obtained exclusively affected offspring (100% RNAi phenotype) before eventually obtaining 0% phenotype (Fig. 3D: Experiments 1, 3a, and 3b). Secondly, RNAi penetrance fluctuates and wanes even when the number of females and egg laying rate are steady (Fig. 3G-H). Thus, while we cannot formally exclude individual differences in reproductive senescence [40], decline in RNAi penetrance was not simply due to death of females in which RNAi was more effective.

### Multiple, independent losses of the dsRNA importer SID-1 in arthropods

For the transit of dsRNA through the mother to the egg, diverse receptor proteins have been implicated in dsRNA cellular uptake and oocyte provisioning. In widening our investigation of the molecular mechanisms of pRNAi, we took a phylogenomic approach to explore the potential relevance of selected receptor proteins in insects. Moreover, our analyses demonstrate a systematic approach for conservation assessments that combines extensive orthology clustering datasets with curation and phylogenetic analysis.

RNAi requires that dsRNA is taken up into the cells of the body, where Dicer acts in the cytosol [4, 5]. The SID-1 protein is a transmembrane importer of long dsRNA and has been a central focus of RNAi research. First characterized in *C. elegans* [41], it is one of several functionally related proteins whose absence causes a systemic RNA interference deficient (SID) phenotype (reviewed in [42, 43]). Conservation of SID-1 is in fact notably variable across insect species, with homologues somewhat agnostically referred to as SID-1-like (SIL) or SID-1-related (Sir) [12]. Nonetheless, ever since early recognition of SID-1 homologues in *Tribolium* and vertebrates [41], it is routinely sought when characterizing RNAi components in new transcriptomes and genomes (see Discussion).

In the last five years the substantial increase in available genomic resources, particularly for the wider diversity of insects [44], enables a more systematic approach based on official gene set (OGS) data from sequenced genomes. Here, we make use of the latest version of the orthology clustering database OrthoDB to survey 148 insect species, embedded in the evolutionary framework of 448 metazoan animal species ([45], Fig. 4: cladogram).

**Fig. 4.**
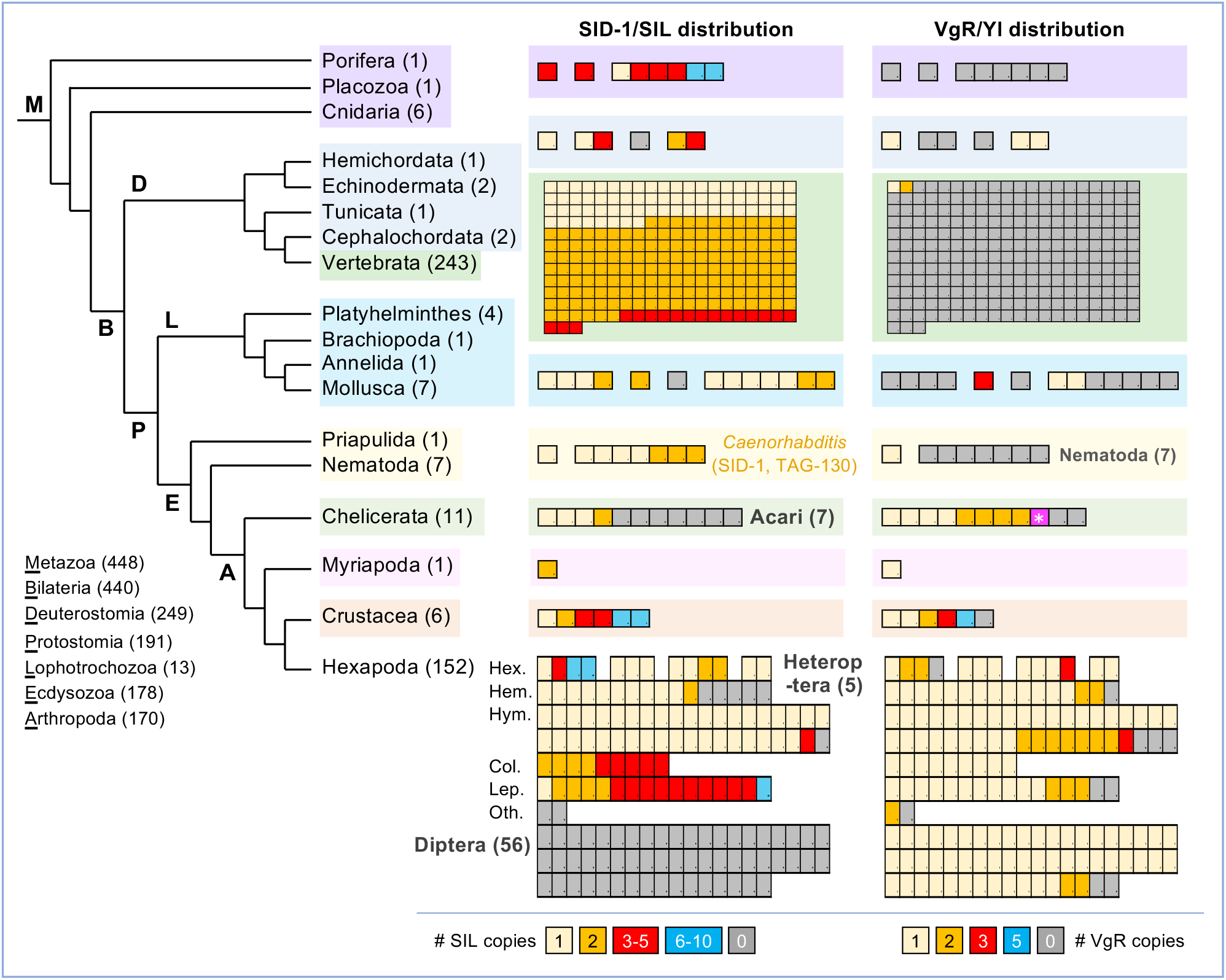
Visualization of metazoan orthology clustering reveals macroevolutionary patterns of protein conservation and lineage-specific losses. Taxonomic distribution and copy number of the SID-1/SIL and VgR transmembrane receptor proteins, representing all metazoan animal species in OrthoDB v10.1, with species numbers stated parenthetically. Phylogenetic relationships are based on [87, 88]. Protein distributions are shown with one box per species, ordered sequentially by copy number, with the color code indicated in the legend for each gene. Notable lineage-specific absences are indicated in bold grey text. For one mite species (Acari), a VgR protein was only included in the wider metazoan orthology group, but this species did not have a VgR protein based on orthology clustering of Arthropoda only (magenta with white asterisk). No other presence/absence results differed across the Insecta, Hexapoda, Arthropoda, and Metazoa clustering analyses. For minor changes in copy number across clustering analyses, the value reported here is based on the most taxonomically restricted analysis (see Methods). Hexapoda taxonomic abbreviations and species counts: Hex.: Non-insect Hexapoda (4), Palaeoptera (3), Polyneoptera (4), Non-hemipteran Paraneoptera (2); Hem.: Hemiptera (16); Hym.: Hymenoptera (40); Col.: Coleoptera (9); Lep.: Lepidoptera (16); Oth.: other Holometabola: Strepsiptera (1), Trichoptera (1). Vertebrate SID-1 proteins are mostly multi-copy, with single orthologues in ray-finned fishes (Actinopterygii), some orders of birds (Pelecaniformes, Gruiformes), and the platypus.

Our assessments of orthology group membership at the hierarchical taxonomic levels of Insecta, Hexapoda, Arthropoda, and Metazoa substantially extend previous observations on the distribution of SID-1 (Fig. 4: “SID-1/SIL distribution”; see Methods and Discussion). Across the Metazoa, SID-1 proteins are present in 375 species, with multiple copies found in 235 of these species. As previously documented with limited sampling [12], we find lineage-wide copy number increases within each of the sarcopterygian vertebrates (the lobe-finned fishes clade, including mammals), Coleoptera (beetles), and Lepidoptera (moths and butterflies). This includes the three SIL proteins originally characterized in *Tribolium* [12]. At the same time, SID-1 is absent from all 56 species of Diptera and 7 Acari species, augmenting previous reports [46, 47]. Furthermore, we newly report the complete absence of SID-1 homologues in an additional, independent lineage: the Heteroptera (true bugs) within the insect order Hemiptera (Fig. 4). To corroborate these evolutionary changes, we further scrutinized OGS, genome assembly, and transcriptome analysis data.

Orthology clustering indicates the lineage-specific loss of SID-1 within the Hemiptera based on its absence in five Heteroptera and presence in eleven outgroup species (formerly the paraphyletic “Homoptera”, including aphids, psyllids, and planthoppers; Fig. 4). To augment species sampling, we compiled recently published results and conducted BLAST investigations of assembled genomes (see Methods), nearly doubling the number of species investigated (Fig. 5A). Importantly, directly interrogating genome assemblies overcomes limitations of OGS gene model predictions [48, 49]. Our tBLASTn searches with diverse SIL orthologue queries did not detect any heteropteran or dipteran sequences but did recover all SIL proteins in other insects (Fig. 5B). Thus, loss of SID-1/SIL spans the four major infraorders of Heteroptera (10 species) compared to its retention in other Hemiptera (present in 15 species, with absences confined to three taxonomically scattered species with limited transcriptomic evidence; Fig. 5A).

**Figure 5.**
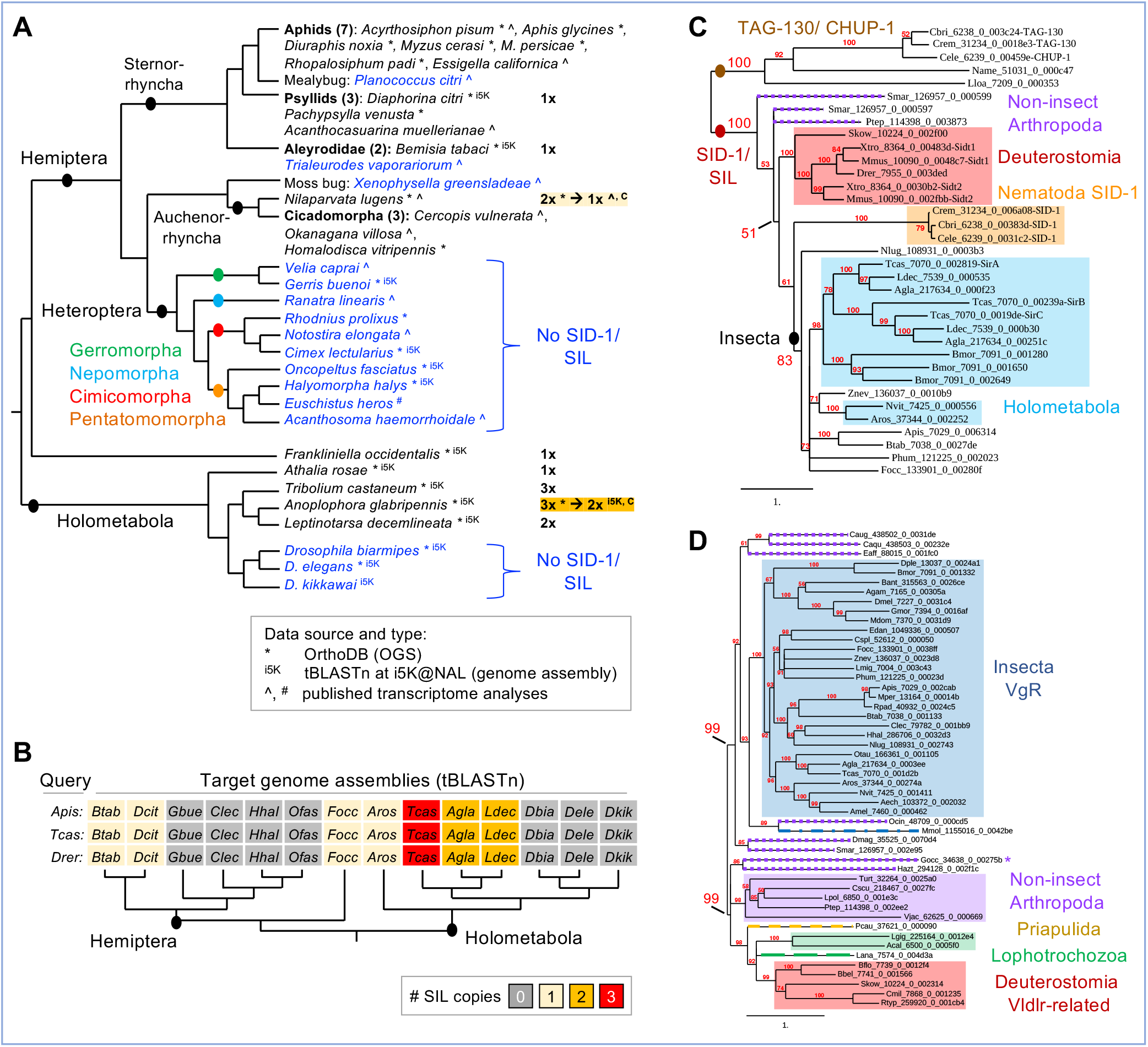
Curation, BLAST, and phylogenetics confirm and refine orthology clustering assessments of SID-1 and VgR distributions. **(A)** Detailed evaluation of genomic resources for Hemiptera and selected outgroups supports the lineage-specific loss of SID-1 in the Heteroptera: species in blue text lack SID-1. Data types and sources are indicated in the legend, including recent transcriptomes (^:[46]; ^#^: [7]), genome assemblies (^i5K^: [78]), and OGS collections at OrthoDB (*: [45]). Phylogenetic relationships after [88-90]. For two species (*Nilaparvata lugens* and *Anoplophora glabripennis*), follow-up curation (“C”) reduced SID-1 copy number compared to the OrthoDB assessment, as indicated (see Methods). **(B)** Selected subset of 14 species from (A) that were further interrogated by direct tBLASTn searching of the genome assembly. Each of the three orthologous query proteins from *A. pisum, T. castaneum* (SirA), and *Danio rerio* produced identical outcomes for copy number. **(C-D)** Maximum likelihood whole-protein phylogenies of SID-1 homologues based on 35 proteins from 23 species (C) and VgR/Vldlr homologues based on 50 proteins from 50 species (D). The branch length unit representing substitutions per site. All nodes have ≥50% support (enlarged labels for selected nodes). Shaded boxes indicate clades of interest, as labeled in the figure, with dashed colored lines for paraphyletic protein members. For the VgR/Vldlr tree, the protein marked with an asterisk (*) represents the chelicerate species that was only included in the Metazoa, but not the Arthropoda, orthology clustering analysis (see Fig. 4).

Even with more extensive species sampling than was previously possible [12, 46], some of the same phylogenetic ambiguities of SIL proteins remain (Figs. 5C, S1). Within *Caenorhabditis* nematodes, SID-1 has high sequence similarity to the functionally unrelated TAG-130/CHUP-1 protein (Figs. 4, 5C; [12]). Our phylogenies are generally robust for topology within clades for the insects and the deuterostomes, but the long-branch nematode proteins are unstable. Two nematode species with single-copy orthologues have particularly long branches and tend to show affinity with *Caenorhabditis* TAG-130. However, the recovery of well supported clades for each of SID-1 and TAG-130 in *Caenorhabditis* species is inconsistent (Fig. S1A-C). In our phylogeny with broad species sampling, all arthropod and deuterostome proteins show greater affinity to nematode SID-1 (Fig. 5C). Lineage-specific duplications appear ancestral, with a single duplication at the base of the sarcopterygian vertebrates and the beetles, and two at the base of the Lepidoptera (Figs. 5C, S1B,D, but with unstable topology for *Tribolium* SirB). The Hymenoptera (wasps, bees) are an outgroup to other Holometabola, yet their single-copy SIL orthologues group elsewhere (Figs. 5C, S1D). Overall, sequence-based assessments of SID-1/SIL conservation are complicated by lineage-specific duplications and rates of sequence evolution, even before its functional relevance for RNAi in insects is considered (see Discussion).

### Maternal provisioning uses distinct receptor proteins in insects and nematodes

An alternative, long-recognized mechanism of dsRNA cellular uptake is endocytosis, for which core genes are widely conserved as standard eukaryotic cellular machinery [4, 42]. Receptor-mediated endocytosis also supports maternal provisioning of oocytes, and it has been proposed for invertebrates that yolk proteins (vitellogenins) and dsRNA may share a common import mechanism [22, 23]. We thus applied our phylogenomic approach to determine conservation of the vitellogenin receptor (VgR), known as Yolkless (Yl) in *Drosophila* (Figs. 4, 5D).

We find a fundamentally different distribution for VgR compared to SID-1 (Fig. 4: “VgR/Yl distribution”). Whereas SID-1 had orthology group members extending to the non-bilaterian Metazoa, VgR is essentially restricted to the Ecdysozoa, excluding the Nematoda. Secondly, whereas there is evidence for multiple VgR proteins in other arthropod groups, this protein is predominantly single-copy throughout the insects, including the Heteroptera and Diptera, and the Coleoptera and Lepidoptera – which lost or duplicated SID-1, respectively. Unlike SID-1, for VgR there are also scattered single-species absences throughout the hexapod orders.

Curiously, two species are the sole exception to the complete absence of vertebrate protein members from the metazoan VgR orthology group (Fig. 4). Our phylogenetic appraisal centered on this anomaly. We obtain two strongly supported clades containing either insect VgR or the deuterostome proteins, with a paraphyletic splitting of non-insect arthropod proteins between these two clades (Fig. 5D). Tracking the vertebrate proteins into the more taxonomically restricted Vertebrata orthology group revealed that these proteins are divergent members of the Very Low-Density Lipoprotein Receptor (Vldlr) proteins, which are conserved in all 243 vertebrate species. In summary, the broad distribution patterns suggested by orthology clustering alone are valid, with our follow-up analyses refining this to strongly support a hexapod-specific origin of VgR. Thus, for the purposes maternal provisioning of oocytes, nematodes and insects rely on distinct receptors.

## DISCUSSION

Our tripartite investigation of the molecular mechanism of pRNAi in *Tribolium* combines (1) an RT-qPCR strategy that detects dsRNA transmitted to the egg, (2) time course assays that show months-long persistence of pRNAi under different parameters, and (3) a phylogenomics profiling approach for appraisal of candidate genes’ taxonomic distributions. Our surprising empirical observations can inform experimental design for developmental genetics studies and targeting strategies for RNAi-based pest management applications. Furthermore, we highlight several key steps at which the cellular mechanism of dsRNA transport remains unresolved, despite highly effective use of RNAi in insects for decades [1, 2, 5, 6, 15].

### Amplicon design and developmental staging determine measured knockdown efficiency

We show that comparison of RT-qPCR results between nested and semi-nested amplicons is a robust method for detection of maternally transmitted dsRNA in eggs (Figs. 1-2). Complementing short-term tracking of fluorescently labeled dsRNA [6, 22, 23], our method detects dsRNA throughout embryogenesis. On the other hand, use of a nested amplicon alone may lead to underestimation of knockdown efficiency, or even to erroneous interpretations of target gene overexpression, depending on endogenous expression levels. Awareness of these features can be applied to tracking dsRNA and to mitigate against unwanted dsRNA detection in single-amplicon assays.

For a gene of interest, primer design may be constrained such that an RT-qPCR amplicon is nested within the dsRNA region. To design intron-spanning primers for short, efficient amplicon sizes [28, 29], while also avoiding conserved coding sequence regions that could cause off-target effects [15, 20], both RT-qPCR and dsRNA primers may target the same region. Small genes with few introns are particularly constrained, such as *Tc-zen1* (Fig 1A: Fragment 3 with respect to the short dsRNA that avoids the homeobox, as in [20]). Secondly, for efficient screening of both expression and function, a single longer amplicon may serve as template for both *in situ* hybridization, where probe sensitivity correlates with sequence length [50], and for RNAi, where longer dsRNA is more effective [10]: this is the case with the long dsRNA for *Tc-zen1* examined here (Fig 1A, [25]).

We find that nested amplicons underestimate true knockdown strength by 5-20% compared to measurements with semi-nested amplicons that only detect endogenous transcript (Figs. 1B, 2A-B). Yet in previous work we consistently obtained strong knockdown validation with a nested amplicon, to 10% of wild type levels ([20]: Fragment 3 and the short dsRNA, Fig. 1A). A key factor was tight developmental staging that targeted peak endogenous expression. Broad sampling beyond the peak expression window effectively dilutes the detection of wild type endogenous transcript levels as the baseline against which RNAi samples are compared. This can substantially alter calculations of knockdown efficiency (Fig. 6), whether using nested or semi-nested amplicons. Thus, staging precision is critical for accurate detection of knockdown efficiency, and this can largely overcome the underestimation effect of using a nested amplicon.

**Figure 6.**
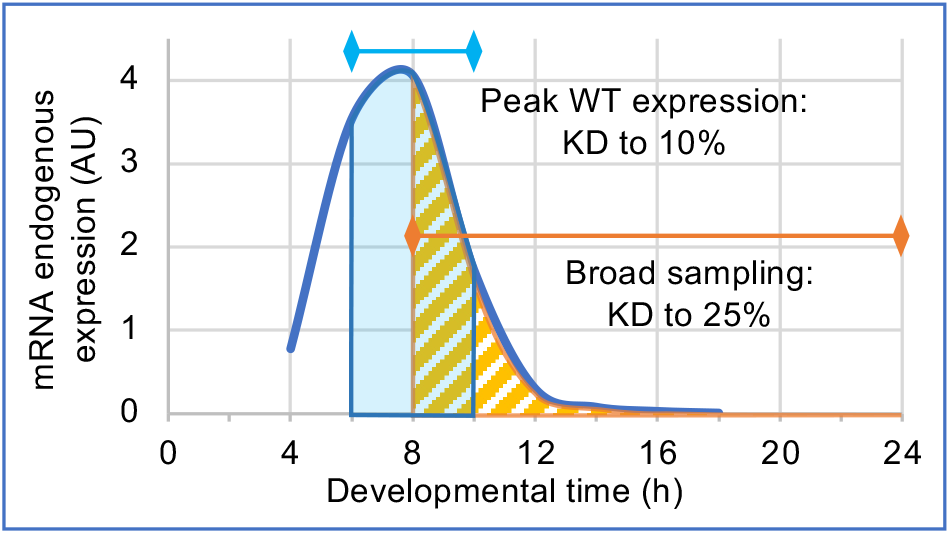
Tighter developmental staging mitigates underestimation of RNAi knockdown when assayed with a nested qPCR amplicon. This schematized representation based on empirical data for *Tc-zen1* illustrates how the time window assayed by RT-qPCR compares to the time course of endogenous expression [20], and in turn how this affects the apparent efficiency of RNAi knockdown. Even with a nested amplicon, assays that strictly target the time window of peak endogenous expression confirm strong knockdown to 10% of wild type levels (blue: based on use of Fragment 3 depicted in Fig. 1A, [20]). In contrast, broad sampling that includes periods of low endogenous expression are more susceptible to underestimation of knockdown (calculated as 25% of wild type levels), due to unmasked detection of dsRNA with a nested amplicon (orange: based on Fragment 2, data in Fig. 1B). Equally, for *Tc-chs1* we obtained two-fold variation in calculated knockdown level from different developmental stages of the same experiment, with either nested or semi-nested amplicons (Fig. 2A).

Measured expression levels are also affected by sequence-specific features. We most strongly detected dsRNA for medial regions of the *Tc-zen1* molecule, with a five-fold decrease towards the 3’ and 5’ ends (Fig. 1C). We therefore speculated that a dsRNA degradation mechanism may lead to progressive loss of detection from both termini. However, a 5’ terminal amplicon detected stable dsRNA levels throughout embryogenesis (Fig. 1B: latter three stages with Fragment 2), arguing for alternative explanations. On further scrutiny, we find that minor differences in amplicon length strongly negatively correlate with amplification efficiency (Fig. S2, [29]). Also, despite primer specificity, we cannot exclude the possibility that our medial amplicon (Fragment 4) may weakly detect the homeobox of the closely related paralogue *Tc-zen2* [20, 51].

Overall, it is striking that long dsRNA is stable *in vivo* in insect eggs, and our nested amplicon strategy offers new opportunities for dsRNA quantification and long-term tracking.

### pRNAi application in relation to knockdown persistence and female fecundity

While confirming that pRNAi wanes within individual females (Fig. 3, [1, 3, 36]), unexpectedly we find that this only occurs after strong knockdown for nearly nine weeks – far longer than was previously shown or assumed. Early research in *Tribolium* reported substantial waning by three weeks after injection and complete cessation of knockdown by five weeks [1]. Accordingly, developmental genetics research generally examines eggs in the first 4-20 days after injection (*e*.*g*., [25, 36, 52]), although ≥90% phenotype penetrance for up to 4.5 weeks has been shown [11]. Differing knockdown durations may reflect differences in injection age (pupal or adult), gene-specific RNAi efficiency [20, 36], and strain-specific rates of waning (Fig. 3D). More generally, our results demonstrate the potential for high-efficiency, persistent pRNAi-mediated knockdown, even after a single instance of dsRNA delivery.

It is also surprising that after 50 days there was an abrupt increase in fecundity in both beetle strains used in this study (Fig. 3F). It was in this time window of intermediate female age (50-100 days) that we obtained fecundity levels comparable to previous reports, which examined the first two months in a third strain (San Bernardino strain: [1, 53]).

These observations highlight within-species variation in the onset and duration of peak fecundity and the rate of RNAi waning. Extrapolation from our study under laboratory conditions (at 30 °C) could also imply longer durations of peak fecundity in natural environments, for slower life cycles at cooler ambient temperatures [e.g., 54]. These factors should be taken into account when planning seasonal management of agricultural pest species by RNAi [5, 6].

### Genomic loss and ambiguous homology of SID-1 emphasizes its minimal relevance for RNAi outside of nematodes

The SID-1 channel protein has been part of the standard repertoire of RNAi-associated cellular machinery in surveys of transcriptomes and genomes (*e*.*g*., [7, 12, 41, 46]). However, our metazoan-wide appraisal confirms multiple lineage-specific losses of SIL from arthropod genomes (Figs. 4-5) and that this protein family encompasses homology across SID-1 and TAG-130/CHUP-1 proteins (Figs. 5, S1). This strengthens a cumulative body of evidence in insects for ambiguous homology and limited functional relevance of SIL for RNAi [4, 5, 12, 42, 46].

The loss of SIL proteins is far more pervasive than previously recognized. Among the chelicerates, its absence in the Acari (mites and ticks) contrasts with retention in spiders and scorpions (Fig. 4, [47]). Its absence in flies [12, 41] may reflect ancestral genomic loss in the wider lineage Antliophora (Diptera, Mecoptera, and Siphonaptera, [46]). For other lineages, reports on single or few species noted anecdotal absences, including in the Heteroptera [7, 42, 46]. A recent review of RNAi specifically in the Hemiptera thus only reported general conservation of SID-1/SIL proteins in this order [6], without recognizing its wholesale absence in the true bugs (Figs. 4-5). Species sampling to date also supports SIL loss in the Trichoptera (Fig. 4 and [46]: 3 species), which may be further borne out as insect genomic resources continue to grow.

Multiple SIL losses in arthropods may seem surprising compared to its vertebrate-wide retention and the fact that nematodes and arthropods are more closely related as fellow Ecdysozoa (Fig. 4). This could suggest a higher rate of evolutionary divergence in arthropods against a backdrop of bilaterian-wide conservation. In fact, vertebrate protein homology suffers from the same ambiguities as analyses with arthropod proteins (Fig. S1). Vertebrate Sidt proteins show greater sequence similarity in certain functional motifs with TAG-130/CHUP-1 proteins, recognized for their role in <u>ch</u>olesterol <u>up</u>take [55]. Furthermore, recent cell culture work suggests that prior evidence for dsRNA uptake by Sidt/CHUP-1 may have detected a secondary consequence of dsRNA association with imported cholesterol [56], calling Sidt molecular function into question. Overall, this is conceptually similar to the macroevolutionary “functional lability” and repeated lineage-specific loss of RNA-dependent RNA polymerases (RdRPs, [57]), another component of systemic RNAi in some species (see below).

In *C. elegans*, SID-1 is required for the systemic spread of RNAi within somatic tissues and the pRNAi effect in offspring [41]. Yet, despite the absence of any SID-1/SIL protein, the Heteroptera are highly sensitive to RNAi (reviewed in [58]). Knockdown is effective and systemic within the bodies of individual heteropteran nymphs [59]. pRNAi can achieve complete phenotypic knockdown in >95% of progeny for at least three weeks [60].

Thus, just like other nematode SID proteins [4, 5, 43], SID-1 should be retired from general inclusion among the insect RNAi repertoire.

### The power of orthology clustering, in context

As discussed, some of our key insights into the taxonomic distribution of SID-1 were already documented on an anecdotal level in a range of published studies, but they had not been integrated. We show that metazoan-wide orthology clustering [45] combined with taxonomically-informed visualization (Fig. 4) can reveal previously unappreciated macroevolutionary patterns of protein origin, conservation, duplication, and loss across disparate lineages such as insects and vertebrates. With corroboration from additional lines of evidence including protein member curation, genome searches, phylogenetics, and literature surveys (Fig. 5), this is a powerful approach.

Such rapid phylogenomic profiling (Fig. 4) could be widely applied to whole suites of proteins, providing criteria for candidate gene selection alongside standard gene ontology (GO) features such as molecular function (transmembrane receptor) or biological process (receptor-mediated endocytosis). And, while our focus is the insects in general, visualization can be customized for other taxa of interest (*e*.*g*., Vertebrata, Hymenoptera), particularly as the number and diversity of sequenced genomes increases.

Orthology clustering across distantly related species requires care. Whereas wholesale loss or duplication in a clade is convincing, taxonomically scattered copy number changes may reflect genuine evolutionary change in undersampled lineages or limitations in individual species’ data quality. Manual curation is necessary to eliminate redundant isoforms, which inflate copy number (Fig. 5A), and incomplete or suspiciously large and divergent proteins, which often reflect inaccurate gene model annotation [48, 49] and can skew phylogenetic analysis (see Methods). Secondly, each taxonomic level of orthology clustering is an independent analysis. At wider taxonomic levels, groups of single-copy orthologues often gain divergent within-species homologues and appear multi-copy due to greater sequence divergence between homologues in distantly related species. The inclusion of divergent vertebrate Vldlr proteins within the metazoan-level orthology group for VgR exemplifies this (Fig. 4). The challenge of reconciling clustering analyses across taxonomic levels is a known, but perhaps not widely appreciated, issue [61]. Clarification of orthology is possible by prioritizing taxonomically restricted clustering results and then progressively adding wider taxa (*e*.*g*., from Insecta to Metazoa, Fig. 4), supported by phylogenetic analysis (Fig. 5). However, the SID-1 and TAG-130/CHUP-1 proteins are particularly recalcitrant, forming a single orthology group even within the Nematoda alone.

### How can pRNAi persistence be reconciled with dsRNA cellular processing and maternal transmission?

Our unexpected finding that the long dsRNA molecule is maternally transmitted into eggs, thereby depleting maternal dsRNA levels, is difficult to reconcile with pRNAi persistence for months (Figs. 1-3). We also find limitations in attributing dsRNA cellular transmission to specific import proteins (Figs. 4-5). Furthermore, biochemical, physiological, and cellular studies on dsRNA processing highlight where dsRNA is *not* located, rather than how it is delivered to Dicer to trigger RNAi. To conclude, we discuss how our observations fit into the wider framework of outstanding major questions on systemic parental RNAi insects (Fig. 7).

**Figure 7.**
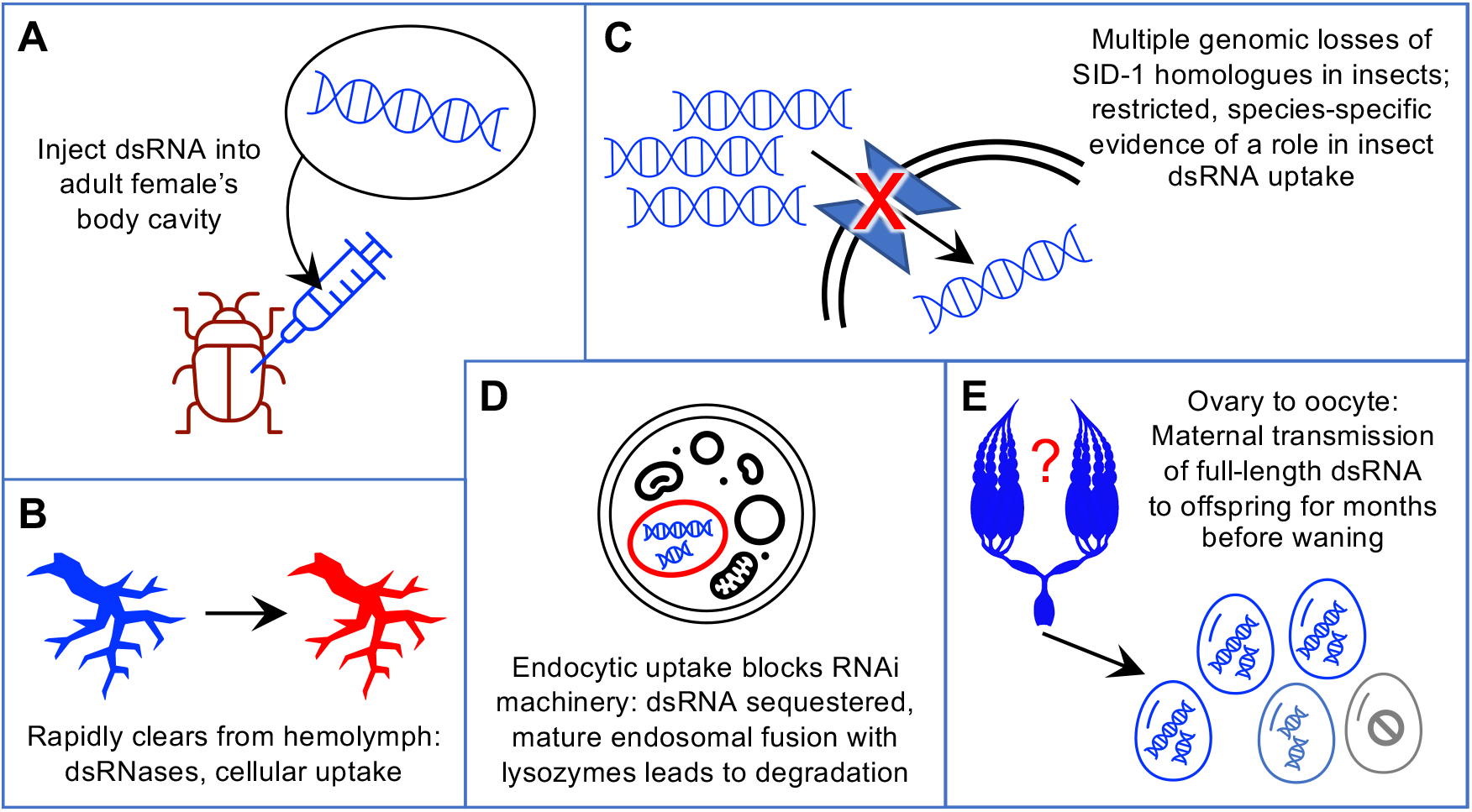
Unresolved features of systemic parental RNAi. Where is the dsRNA stored long-term in the mother without degradation and with continuous transmission to eggs? Cartoons represent the progression of dsRNA from initial injection **(A)**, through the mother’s tissues **(B)** and cells **(C**,**D)**, to the oocytes **(E)**. Presence of dsRNA is represented in blue, with specific cell- and tissue-scale challenges to its transmission shown in red, and with final waning of pRNAi indicated by pale blue and grey. Clip art images reproduced and modified from Microsoft PowerPoint 2021, v. 16.52.; ovary silhouette based on image at https://cronodon.com/BioTech/Insect_Reproduction.html.

Upon injection into the female’s body cavity (Fig. 7A), dsRNA spreads throughout the circulatory system. However, it rapidly clears – on the scale of minutes to hours – from the hemolymph due to cellular uptake and degradation (Fig. 7B, [21, 23, 62]). In *Tribolium*, substantial activity of endogenous dsRNases is documented in the gut and implicated in the hemolymph [63]. Also, the ovary represents just one organ in the female body in which dsRNA uptake occurs. In effect, the germline competes with other cell types for dsRNA. Particularly when it is distal to the site of dsRNA injection, it may be less sensitive or even refractory to RNAi [9, 23]. Injection of dsRNA for pRNAi is highly effective in practice, but not without limitations.

Second, the dsRNA received by the insect ovary represents a non-renewable resource. In this and other studies, pRNAi is achieved after a single injection, providing a finite number of dsRNA molecules. That starting pool is amplified by RdRPs in plants, nematodes, and possibly fungi [57, 64-66]. This property can be exploited *in planta* for sustained delivery of non-endogenous transcripts in RNAi-based pest control [64]. However, there is no evidence to date for dsRNA amplification in insects (reviewed in [57, 63]). Also, amplification in other species generally or exclusively involves siRNA synthesis [64-66], which contrasts with our detection of ≥100-bp RT-qPCR amplicons spanning full-length long dsRNAs (Figs. 1-2).

Next, there are uncertainties as to how cellular uptake of long dsRNA is accomplished (Fig. 7C). In principle dsRNA could be shuttled into the oocyte after uptake by the nurse cells or the follicular epithelium, or it could be directly imported by the oocyte during patency, when intercellular openings in the follicular epithelium confer direct access to the hemolymph. However, neither SID-1 for cellular uptake (discussed above) nor VgR for oocyte endocytosis seems to be the effector. In *C. elegans*, co-accumulation of dsRNA and vitellogenin in oocytes suggested a common import mechanism for these molecules [22].

However, the VgR receptor is hexapod-specific (Figs. 4-5), arguing against a conserved mechanism associated with invertebrate vitellogenin transport. Furthermore, trials with labeled dsRNA revealed its exclusion from oocytes during vitellogenesis [23]. On the other hand, SID-1 and VgR are two candidates among many potential receptor proteins. Clathrin-dependent endocytosis is required for within-individual larval RNAi in *Tribolium* [24], and such mechanisms may also be applicable for pRNAi.

More generally, endocytosis has long been recognized as a potential mechanism for dsRNA uptake, but it has its own cellular challenges (Fig. 7D, reviewed in [4, 5, 42]). First, if dsRNA is sequestered within an endosome, it is inaccessible for processing by Dicer in the cytosol, and the mechanism of selective endosomal escape of dsRNA is unknown. Species-specific levels of dsRNA sequestration have been correlated with susceptibility to RNAi [5]. Second, endosome maturation culminates in fusion with a lysosome, targeting all contents for degradation [4]. Thus, endosomes do not seem suitable as long-term, slow-release reservoirs for pRNAi. Beetles including *Tribolium* appear to have low levels of endosomal sequestration, but those studies were performed in larvae [reviewed in 5]. Further investigation of maternal reproductive tissues may reveal alternative, germline-specific mechanisms of dsRNA retention and cell-to-cell transmission. This would be fully consistent with the growing body of evidence for the tissue-specific as well as stage-specific nature of RNAi (*e*.*g*., discussed in [9, 23, 67]).

Finally, dsRNA’s journey from maternal injection through successful embryonic knockdown requires two levels of maternal transmission (Fig. 7E). After dsRNA is delivered into the oocyte, cellular uptake must happen again: when dsRNA within the yolky oocyte is taken up by the embryonic cells, where knockdown is finally achieved. As maternal injection can lead to deposition of labeled oligonucleotides in the yolk without embryonic uptake [68], this step also cannot be taken for granted. In summary, while we continue to successfully use pRNAi for developmental genetics research and in devising new and improved strategies for pest management, there remain many aspects of dsRNA transport and systemic propagation that await explanation.

## MATERIALS AND METHODS

### *Tribolium castaneum* (Herbst) stocks and genomic resources

All beetle stocks were kept under standard culturing conditions [13] at 30°C, 50 ± 10% RH. The lines used for the RT-qPCR assays were San Bernardino (SB) wild type [13] and nuclear GFP (nGFP) [33]. For the RNAi penetrance time course experiments, Strain 1 was a heterozygous cross of the enhancer trap lines G04609 (females; [35]) and HC079 (males; [30]), both in the *pearl* white-eyed mutant background [69]; Strain 2 was the LifeAct-GFP line, in a rescued *vermillion white* background [70].

Sequence data for the target genes in this study are based on the latest genome assembly and official gene set (OGS3, [71]): *Tc-zen1* (TC000921, [20, 26]), *Tc-chitin synthase 1* (*Tc-chs1*, TC014634, [27]), *Tc-Ribosomal protein S3* (*Tc-RpS3*, TC008261, [25]), *Tc-germ cell-less* (*Tc-gcl*, TC001571, [39]), and *Tc-tailup* (*Tc-tup*, TC033536, [15, 37]). Details of primers and amplicon sizes are presented in Table S1, also for the transgene *DsRed2* (based on the *piggyBac* mutator construct: GenBank accession EU257621.1).

### Parental RNAi

Parental RNAi was performed as described [25], with dsRNA resuspended in H_2_O and injected at a concentration of approximately 1 µg/µl (range: 900-1100 ng/µl). Beetles were sexed as pupae (distinguished by genital morphology) and allowed to mature to adulthood. Females were anesthetized on ice and dsRNA was injected into the abdomen. Uninjected females served as wild type controls. Gene-specific knockdown phenotypes were confirmed based on published resources for all genes, using the specific assays described below for each of the RT-qPCR and time course experiments. As *Tc-tup* has thus far only been characterized in a high throughput screening analysis [15, 37], we used two non-overlapping fragments (NOFs) of dsRNA in our experiments (NOF1 for Experiments 1 and 2, NOF2 for Experiment 3: see Table S1). We found no quantitative or qualitative phenotypic difference between the non-overlapping fragments.

### RT-qPCR experiments

Embryos were collected over a period of 20 days after injection. Knockdown efficiency was ensured by: manual assessment of serosal cuticle structure (eggshell rigidity) for *Tc-zen1* [11, 20] and *Tc-chs1* [27], detection of fluorescent signal for *dsRed* [34, 35], and by RT-qPCR for all genes. To evaluate *DsRed* knockdown efficiency by fluorescence screening, only larvae were scored to ensure all offspring had successfully completed embryogenesis and were thus old enough to produce strong 3xP3-DsRed signal.

RT-qPCR and data analysis were performed as described, including TRIzol extraction, DNase treatment and gDNA quality control checks, cDNA synthesis, and Fast SYBR Green detection on an Applied Biosystems 7500 Fast cycler (reagents: ThermoFisher Scientific; TURBO DNAfree Kit, Applied Biosystems; SuperScript VILO cDNA Synthesis Kit, Invitrogen; Life Technologies; respectively) [20, 25]. All samples were run in triplicates (technical replicates) with three samples per treatment (biological replicates). *Tc-RpS3* was used as the reference gene, this being established as more stable across embryogenesis as a single reference gene compared to several alternatives with pairs of reference genes or seven other single genes [25]. Raw data were analyzed using LinRegPCR v12.16 [72, 73] and the expression ratio (R) was calculated using the ΔΔCt method, according to the formula:

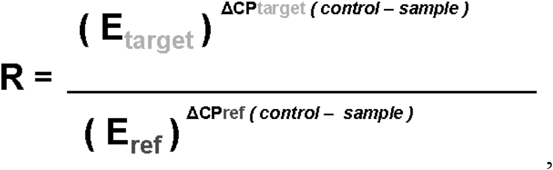

where E is the mean efficiency of the corresponding amplicon as calculated by LinReg and CP is the mean CP of the three technical replicates (after passing quality control in LinReg). The control sample was a pool of all samples (wild type and RNAi; all time points; all biological replicates) of the respective experiment (*i*.*e*., RNAi knockdown of a given gene: *Tc-zen1, Tc-chs1*, or *dsRed*). The % of wild type (WT) was calculated by dividing R_RNAi_ by R_WT_ for the same time point and sample collection date, where both R values are relative to the control sample.

### RNAi penetrance time course experiments

Larval cuticle preparations were used to monitor phenotype penetrance over time after a single injection of dsRNA into the adult female. A cuticle assay is highly effective even with limited embryonic material, which was important in our months-long experiments because female survival and fecundity decline over time [74]. Moreover, *Tc-tup* and *Tc-gcl* provide clear cuticle readouts, whereas RNAi for each of our RT-qPCR target genes can result in non-lethal knockdown that must be analyzed at specific developmental stages (Fig. 2C-F, [27]

Eggs were collected at regular intervals and maintained under standard culturing conditions until a minimum age of ≥4 days after egg lay, to ensure time for larvae to hatch. Larval cuticles were then prepared as described previously [15]. Briefly, eggs and larvae were dechorionated in bleach (VWR # L14709.0F, sodium hypochlorite (11-14% Cl_2_) in aqueous solution), rinsed in tap water, and mounted on slides in 1:1 lactic acid:Hoyer’s solution [75]. Slides were cured overnight at 60 °C to fully clear soft tissues. Slides were then scored under incidental white light on stereomicroscopes, distinguishing six categories: wild type larvae, unhatched wild type (post dorsal closure with no apparent defects, but still at least partially within the vitelline membrane), gene-specific phenotype category 1 (generally a larger body size), gene-specific phenotype category 2 (generally a smaller and less well formed body), non-specific defects, or no larval cuticular material (“empty egg”, indicative of unfertilized eggs or early embryonic lethality). Statistics on penetrance compare wild type with gene-specific knockdown, combining each of the first two categories while for simplicity omitting the latter two, minor categories. The time point of a sample represents the start of the egg collection period (*e*.*g*., data at 3 dpi represent the sample collected 3-4 dpi in Experiment 1, Fig. 3D). Egg collection intervals were extended or pooled to ensure sample sizes of ≥10 offspring per treatment condition for each time point.

Experiments were conducted until three egg collections contained only hatched larvae and the knockdown effect was deemed to have fully waned. Throughout the experiments, dead adult beetles were periodically removed and sexed to note female-specific lethality (males have a darkened cuticular sex patch on the inner/ proximal side of the first leg pair; this is absent in females: https://www.ars.usda.gov/plains-area/mhk/cgahr/spieru/docs/tribolium-stock-maintenance/#sexing [last accessed 15 October 2021]).

To assay females of different ages, adult beetles were maintained continuously under standard culturing conditions at 30 °C until injection. Female age was calculated from the last date when beetles in the experimental cohort were sexed as pupae, reflecting a minor overestimation (≤5 days) relative to eclosion of the adult for some individuals in the cohort. The females used in Experiments 1 and 2 derive from the same cohort and were sexed at the same time.

### Microscopy

Images were acquired on an epifluorescent microscope with structured illumination (Zeiss Axio Imager.Z2 with Apotome.2). Red fluorescence signal in the eyes and ventral nerve cord was used to evaluate *DsRed* RNAi, with green fluorescence from the ubiquitous nGFP signal in this transgenic line serving an internal control. Representative cuticle images were acquired with GFP acquisition settings to detect cuticle autofluorescence, presented as maximum intensity projections from the acquired z-stacks.

### Orthology distribution, BLAST, and phylogenetic evaluations

We examined orthology groups in OrthoDB v. 10.1 [45], comparing the independent orthology clustering analyses at taxonomic levels including Metazoa, Arthropoda, Hexapoda, Insecta, Hemiptera, Coleoptera, Nematoda, and Vertebrata. We noted minor changes in species membership, copy number, and protein ID between the independent orthology clustering analyses conducted at the various taxonomic levels, which is a known issue for orthology clustering [discussed in 61]. In all cases, we used data at the most taxonomically restrictive level (last common ancestor, LCA, level) as the most specific and reliable. For the genes examined here (Fig. 4), orthology clustering was very robust, with only minor differences (*e*.*g*., Fig. 4: asterisk and legend note for VgR).

Curation of protein sequences obtained from orthology groups involved visual inspection of the protein size and sequence in order to remove partial and redundant isoforms. In choosing appropriate protein members of an orthology group for use in phylogenetic analyses, visual inspection of multiple sequence alignments and preliminary trees were used to identify and cull divergent (long branch) proteins and overly long proteins (which may reflect erroneous protein fusion or other model annotation errors such as inclusion of extraneous predicted exons).

Protein sequences were aligned for manual inspection in ClustalW [76], at https://www.genome.jp/tools-bin/clustalw [last accessed 15 October 2021]. Phylogenies were generated at Phylogeny.fr with default settings (alignment with MUSCLE 3.8.31, phylogeny with PhyML 3.1/3.0 aLRT, and tree rendering with TreeDyn 198.3) [77].

Genome assemblies were examined by BLAST, supported by visual inspection of hits with respect to the assembly, gene model predictions, and expression evidence tracks in the Apollo genome browsers, hosted at the i5K@NAL workspace [78]. Species sampling involved a particular focus on the Heteroptera [48, 79-82] and selected species from other orders (Thysanoptera, [83]; Hymenoptera, [84]; Coleoptera, [85, 86]). The genome assembly versions interrogated by tBLASTn are detailed in Table S2.

## SUPPLEMENTARY FILES

**Figure S1.**
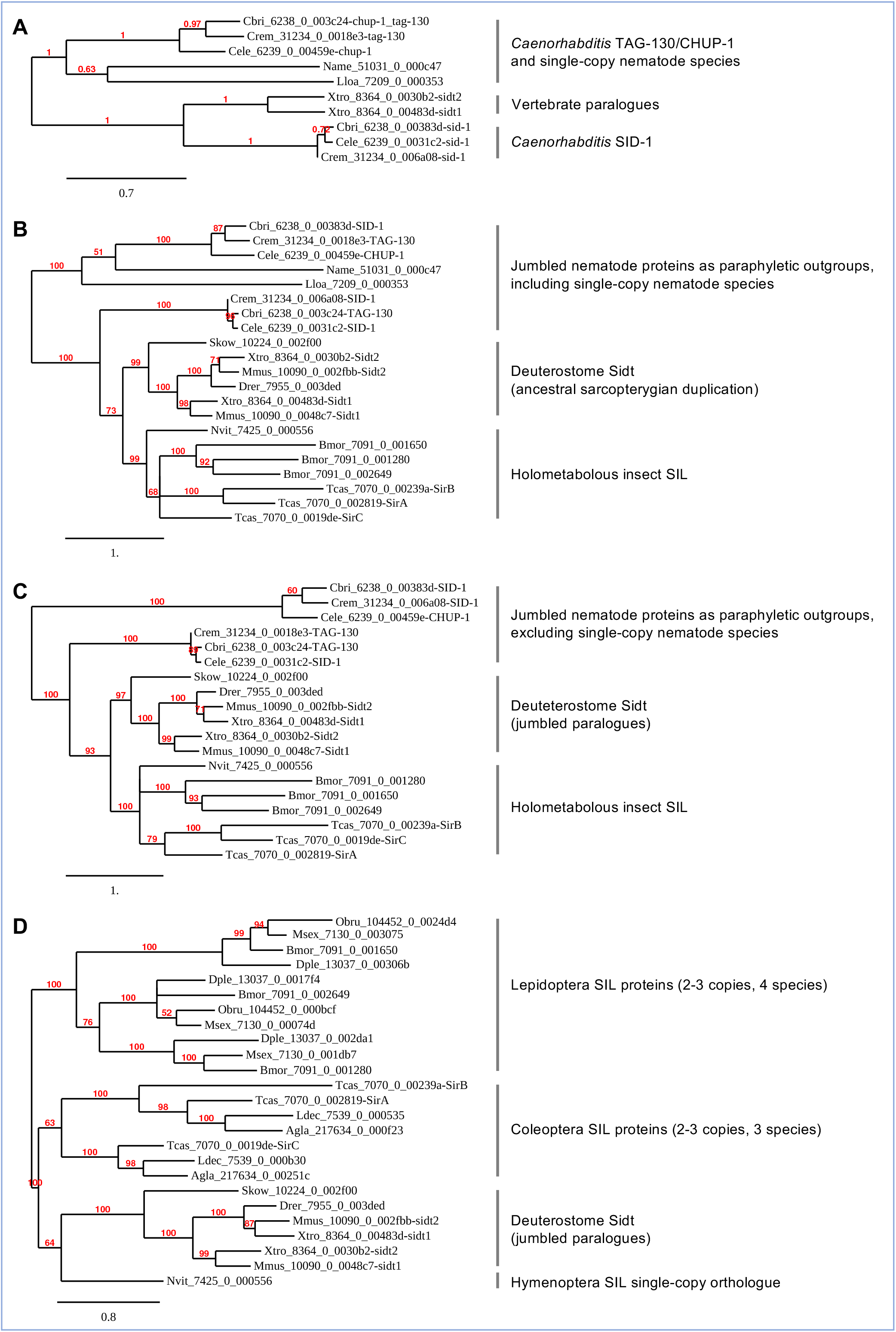
Additional phylogenies with species subsampling for SID-1/SIL proteins. (A-D) Maximum likelihood phylogenies of selected SID-1 homologues. The branch length unit representing substitutions per site. All nodes have ≥50% support. The designation “jumbled” highlights clades that conflate distinct genes (nematode SID-1 with TAG-130/CHUP-1, vertebrate Sidt1 with Sidt2), which did not occur across all trees.

**Figure S2.**
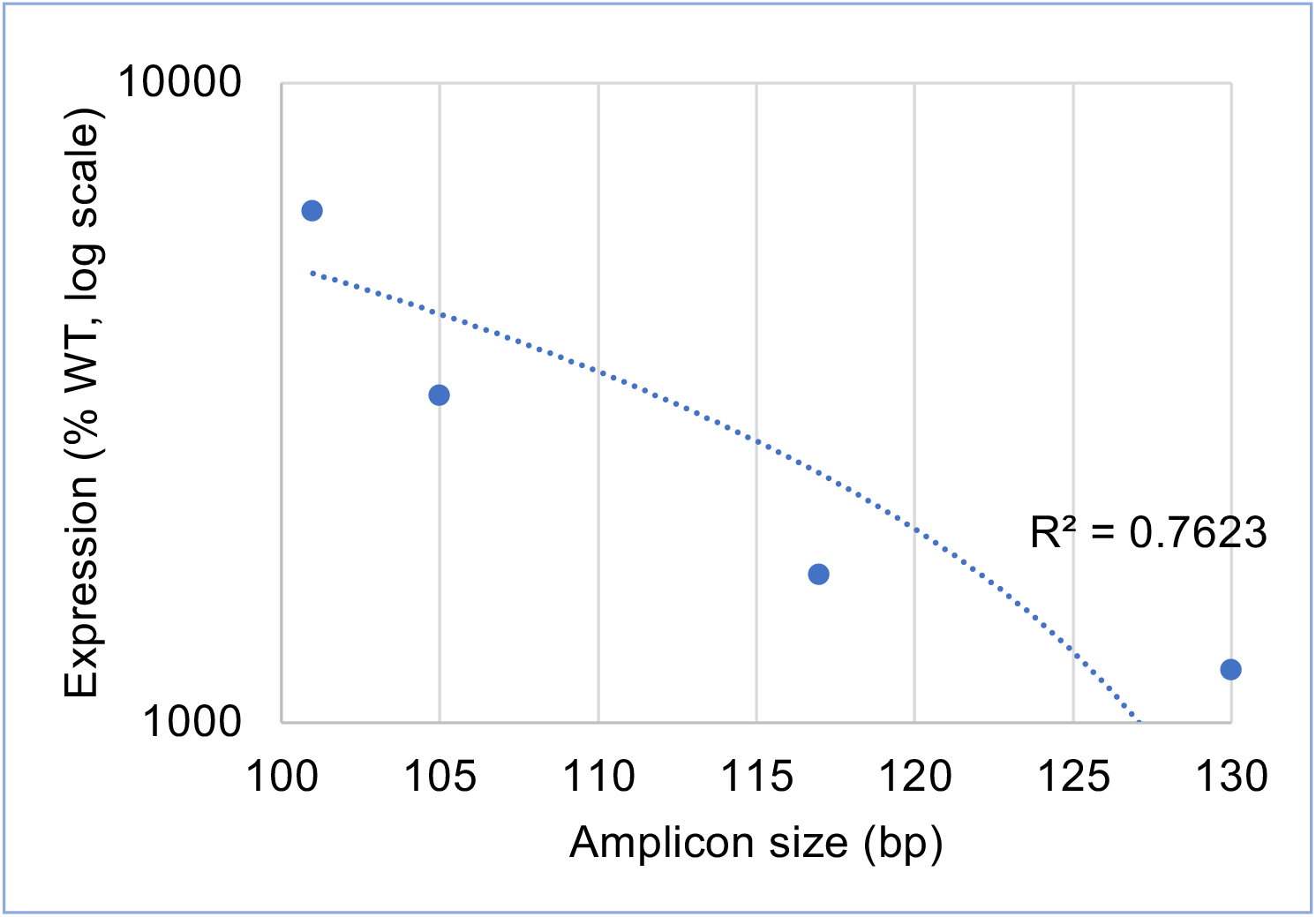
Negative correlation of nested RT-qPCR amplicon length and detection of dsRNA. (expression in excess of WT), assayed for *Tc-zen1* at 16-24 h, as in main text Figure 1C. Fragments 3 and 4 are shorter than Fragments 2 and 5. Logarithmic trendline for mean expression level (% WT) *vs*. amplicon length: R^2^ = 0.76.

**Table S1. Primers used in this study.** Note that primers for RNAi (dsRNA synthesis) also included an adapter sequence, 5’-GGCCGCGG-3’ (forward primers) or 5’-CCCGGGGC-3’ (reverse primers), for subsequent amplification with T7 promoter universal primers (adapters not shown in table). The T7 universal primers are: 5’-universal primer 5’-GAGAATTCTAATACGACTCACTATAGGGCCGCGG-3’, and 3’-universal primer 5’-AGGGATCCTAATACGACTCACTATAGGGCCCGGGGC-3’.

**Table S2. Genome assembly versions queried by BLAST.** These resources were interrogated with tBLASTn queries for selected SID-1 proteins (see main text Figure 5B). Accessed at the i5K@NAL site, most recent access date: 13 October 2021.

## MANUSCRIPT INFORMATION

## Acknowledgments

We thank Stefan Koelzer for generating and processing RNAi samples for RT-qPCR, Siegfried Roth for diverse discussions on insect pRNAi, Robert M. Waterhouse for assistance with and discussions on OrthoDB, Robert M. Waterhouse and Ruixun Wang for detailed and helpful scrutiny of the manuscript, and the Stancliffe Institute for Remote Working for infrastructural support. We also thank Sebastien Santini (CNRS/AMU IGS UMR7256) and the PACA Bioinfo platform (supported by IBISA) for the availability and management of the phylogeny.fr website used for our phylogenetic analyses. Finally, we thank Dominik Stappert for originally designing the long template primer pair for *Tc-zen1* during his doctoral work [74], as this has proven extraordinarily fruitful over the years.

## Author contributions

Conceptualization: TH, KAP; Conducted experiments and analyzed data: TH, KDN, KAP; Primary writing: TH, KAP; Discussion, review, and editing of the manuscript: TH, KDN, KAP.

## Supplementary Figure and Tables

**Table S1.**
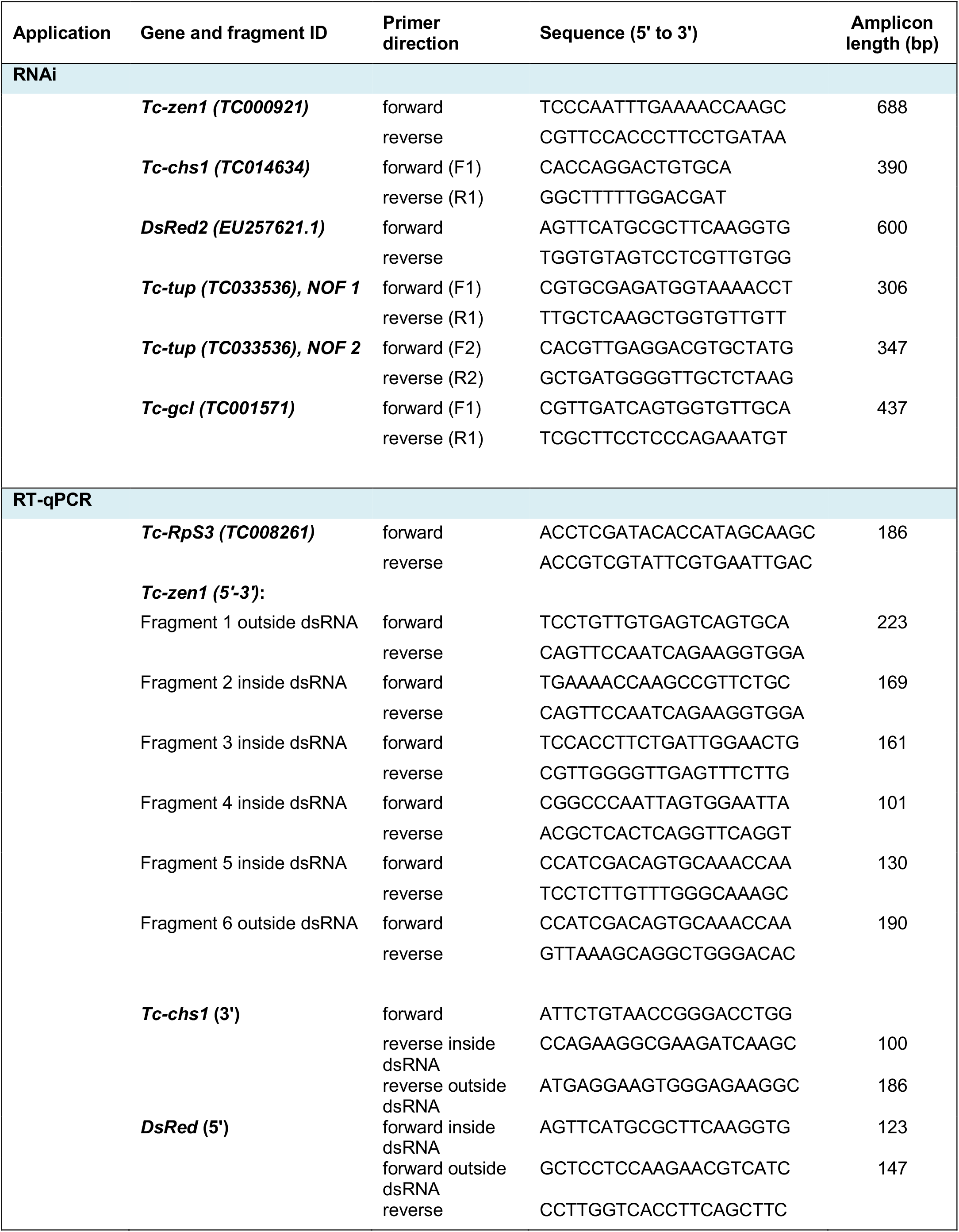
Primers used in the study. Note that primers for RNAi (dsRNA synthesis) also included an adapter sequence, 5’-GGCCGCGG-3’ (forward primers) or 5’-CCCGGGGC-3’ (reverse primers), for subsequent amplification with T7 promoter universal primers (adapters not shown in table). The T7 universal primers are: 5’-universal primer 5’-GAGAATTCTAATACGACTCACTATAGGGCCGCGG-3’, and 3’-universal primer 5’-AGGGATCCTAATACGACTCACTATAGGGCCCGGGGC-3’.

**Table S2.**
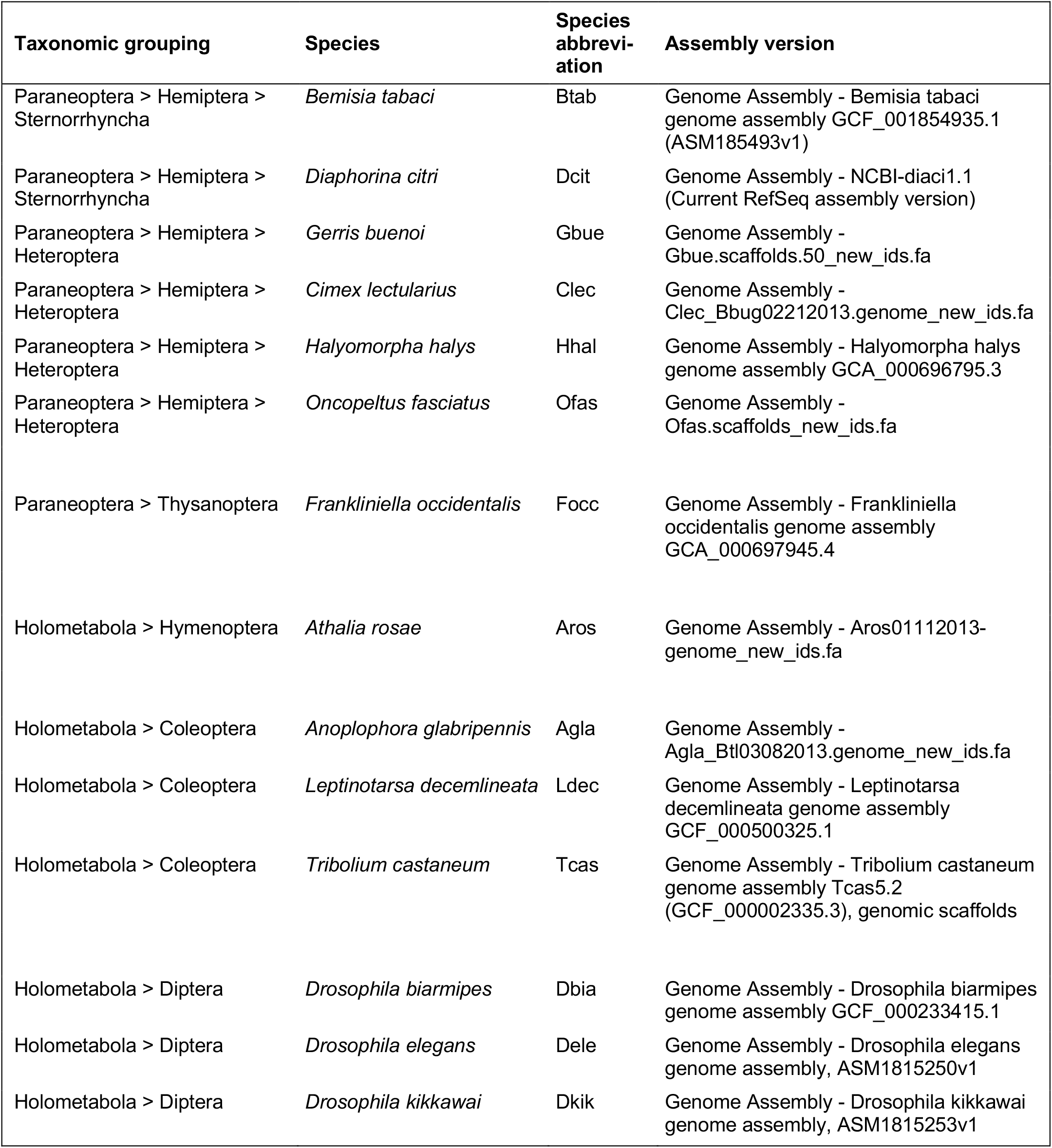
Genome assembly versions queried by BLAST. These resources were interrogated with tBLASTn queries for selected SID-1 proteins (see main text Figure 5B). Accessed at the i5K@NAL site, most recent access date: 13 October 2021.

